# Performances of bioinformatics tools for the analysis of sequencing data of Mycobacterium tuberculosis complex strains

**DOI:** 10.1101/2022.07.05.498825

**Authors:** Pauline Quagliaro, Samira Dziri, Fatma Magdoud El Alaoui, Patrick Saint Louis, Loïc de Pontual, Julie Marin, Etienne Carbonnelle, Typhaine Billard-Pomares

## Abstract

Whole genome sequencing of *Mycobacterium tuberculosis complex* (MTBC) strains is a new and rapidly growing tool to obtain results regarding resistance, virulence factors and phylogeny of the strains. Bioinformatics tools presented as user-friendly and easy to use are available online. The objective of this work was to evaluate the performances of two bioinformatics tools, easily accessible on the internet, for the analysis of sequencing data of MTBC strains.

Two hundred and twenty-seven MTBC strains isolated at the laboratory of the Avicenne Hospital between 2015 and 2021 were sequenced using Illumina®(USA) MiSeq technology. An analysis of the sequencing data was performed using the two tools Mykrobe and PhyResSE. Sequencing quality, resistance or susceptibility status and phylogeny were investigated for each strain. Genotypic resistance results were compared to the results obtained by phenotypic drug susceptibility testing performed in the hospital’s routine laboratory.

Using the PhyResSE tool we found an average coverage of 98% against the reference strain H37Rv and an average depth of 119X. No information on sequencing quality was obtained with the Mykrobe tool. The concordance of each tool with the phenotypic method for determining susceptibility to first-line anti-tuberculosis drugs was 95%. Mykrobe and PhyResSE tools identified resistance to second-line anti-tuberculosis drugs in 5.3% and 5.7% of cases respectively. The sensitivity and specificity of each tool compared to the phenotypic method was respectively 70% and 98% for Mykrobe and 76% and 97% for PhyResSE. Finally, the two tools showed 99.5% agreement in lineage determination.

The Mykrobe and PhyResSE bioinformatics tools were easy to use, fast and efficient. The Mykrobe tool had the advantage of being offline and its interface was more user-friendly. The use of these platforms depends on their accessibility and updating. However, their use is accessible to people not trained in bioinformatics and would allow a complementary approach to phenotypic methods for the study of MTBC strains.

## 1. Introduction

Tuberculosis (TB) is an infectious disease caused by the bacterium *Mycobacterium tuberculosis (Mtb)*. The global population structure of *M. tuberculosis* isolates is classified into seven major lineages, each associated with specific human populations including the Indo-Oceanic (Lineage 1), the East Asian (Lineage 2), the East-African-Indian (Lineage 3), the Euro-American (Lineage 4), the West African-1 (Lineage 5), the West African-2 (Lineage 6) the Ethiopian (Lineage 7) distinct from the *M. bovis* clade (1). The transmission occurs between individuals by dispersion of aerosols, from a contagious patient called bacilliferous (2). Worldwide, TB remains one of the most prevalent infectious diseases and becomes the first leading infectious disease killer (3). Approximately a quarter of the world’s human population has latent TB infection and is at risk of developing TB disease, providing a reservoir of TB for decades (4). The World Health Organization (WHO) estimated 10 million new cases of active TB in 2019 (5). Treatment of TB is based on the four major anti-TB drugs: isoniazid (INH), rifampicin (RMP), ethambutol (EMB), and pyrazinamide (PZA). Short courses of treatment lasting 6 months use the four anti-TB drugs in combination for 2 months, followed by INH and RMP for 4 months (6,7). The emergence of antibiotic resistance in TB strains is a global phenomenon that threatens the WHO’s goal of eradicating TB by 2035 (5). Multidrug-resistant (MDR) strains are defined as resistant to the first-line antibiotics INH and RMP. This resistance is of concern because it significantly increases the risk of treatment failure. Additional resistance to fluoroquinolones (FQ) and at least one additional Group A drug (levofloxacin or moxifloxacin, bedaquiline and linezolid) defines eXtensively Drug Resistance (XDR)(8). The management of patients infected by MDR or XDR strains remain actually long and complex, particularly because of the limited choice of second-line anti-TB drugs (3).

Several methods can be used to detect TB drug resistance. The phenotypic method in solid medium remains the reference method, but it takes at least 3 weeks to obtain results, which leads to a delay in the adaptation of anti-TB treatments (9). The phenotypic method in liquid medium allows to obtain a result more quickly, around 14 days. To reduce this delay, genotyping techniques have been developed, that provide results in a few hours. The use of rapid molecular diagnostic tests based on PCR (Polymerase Chain Reaction) method, such as the Xpert® MTB/RIF assay (Cepheid®)(10) and GenoType MTBDRplus (Hain Lifescience®)(11), have been recommended by the WHO since 2008. These molecular methods make it possible to identify mutations in the target genes of the main anti-TB drugs: the *rpoB* gene involved in rifampicin resistance and the *inhA* and *katG* genes involved in isoniazid resistance for the GenoType MTBDRplus test (Hain Lifescience®)(12). However, these techniques are limited by the number of mutations they can detect (13,14).

The advent of whole genome sequencing (WGS) techniques provides a new approach to the study of *M. tuberculosis complex* (MTBC) strains and plays now a leading role in epidemiologic studies of TB. The complete genome sequence of the *M. tuberculosis* reference strain H37Rv was described in 1998 (15). The genome is approximately 4.41 million base pairs in length and encoded approximately 4000 genes. Using WGS, it is possible to detect resistances that escape molecular techniques. According to recent studies, WGS allows to obtain TB drug resistance profiles on average 9 days earlier than phenotypic tests for first-line anti-TB drugs (16,17). Moreover, WGS not only allows the study of known resistance genes but also the characterization of other loci as predictive of resistance or not (18). Finally, the WGS allows to determine the lineage of each strain and thus to better study the circulation of strains throughout the world.

World health organization considers NGS (Next Generation Sequencing) as “invaluable” to guide diagnosis and treatment, but also for early detection of MDR or XDR *M. tuberculosis* outbreaks (19). Accordingly, several software packages have recently been developed to assist in the analysis of NGS generated sequences. (20–24). They work by detecting pre-defined mutations, mostly single-nucleotide polymorphisms (SNPs), from the reads (sequences of DNA fragments read by the sequencer) or an assembled genome and call the strain resistant whenever one of these mutations is detected. The performances of these software vary from one study to another and according to the tool used. To our knowledge, these softwares are not usually used in routine laboratory in France. The objective of this work was to evaluate the performances of two bioinformatics tools, Mykrobe and PhyResSE, for the determination of resistance and the identification of the lineage of MTBC sequenced strains.

## 2. Material and methods

### Culture and identification of MTBC

Clinical MTBC strains were randomly taken from patients during routine care at Avicenne Hospital to represent clinical French epidemiology of TB. This was a retrospective study and the strains included were all stocked strains. Strains were isolated using Coletsos media (BioRad®, Marnes-la-Coquette, France). Identification of MTBC was performed using Bioline TB Ag MPT64 Rapid test (Abbott®, Chicago, United States) and confirmed after extracting the DNA by using GenoType Hain technology (Biocentric®, France).

### Phenotypic testing

Phenotypic DST was determined for the four first-line drugs using BACTEC™ Mycobacterial Growth Indicator Tube™ (MGIT) 960 (Becton Dickinson®, Sparks, MD, USA) liquid culture system according to the manufacturers’ instructions, at the following critical concentrations: RMP, 1.0 mg/L; INH, 0.1 mg/L; PZA, 100.0 mg/L; EMB, 5.0 mg/L. According to the laboratory’s routine protocol, all results showing phenotypic resistance for MTBC isolates were verified in a second experiment. The phenotypic DST was classically used as the gold standard to calculate sensitivity (prediction of antibiotic resistance), specificity (prediction of antibiotic susceptibility), negative predictive value (NPV) and positive predictive value (PPV) of bioinformatic tools.

### Whole genome sequencing

Genomic DNA was extracted from colonies growing on Coletsos media. Denaturation and DNA extraction from the MTBC strains were performed as previously described (25). DNA quantity was assessed using the Qubit 2.0 fluorometer (Thermo Fisher Scientific®, USA). WGS was performed with Illumina® Technology (MiSeq) using the Nextera XT DNA library preparation kits as instructed by the manufacturer (Illumina®, San Diego, USA).

### Bio informatic tools

Two bioinformatics tools were evaluated in this study. The Phylo-Resistance Search Engine (PhyResSE) (http://PhyResSE.org)(21) is a German online tool for processing MTBC-related bioinformatics data. It has been available for use since 2015 and provides the following information: total number of sequences, percentage in GC, Phred score, depth and coverage to the reference strain H37Rv. The number of variants compared to the reference strain, the lineage and the mutations conferring antibiotic resistance were also obtained with this tool. The resistance for first line antibiotics (Isoniazid, Rifampicin, Pyrazinamide, Ethambutol) and second line antibiotics (Streptomycin, Fluoroquinolone, Kanamycin, Capreomycin, Ethionamide) are detected (26). PhyResSE has compiled a catalog of resistance conferring mutations from the literature and their own laboratory data. For each mutation, a probability of finding the mutation on the liquid antibiotic susceptibility test is given as well as a link to the scientific article highlighting the effect of this mutation. These data are regularly updated. The website is accessible through a standard search engine. A private and secure session is accessed using a 32-character key.

The Mykrobe application (http://mykrobe.com)(27) is an English offline application for processing Mtb-related bioinformatics data. It is available for use since 2019. The information obtained using Mykrobe is the presence of a resistance conferring mutation with its depth and the lineage of the strain. The resistance for first line antibiotics (Isoniazid, Rifampicin, Pyrazinamide, Ethambutol) and second line antibiotics (Streptomycin, Fluoroquinolone, Kanamycin, Capreomycin) are detected (26). No information on the quality of the sequencing is given. The application is accessible on the Internet using a standard search engine and can be easily downloaded.

### Ethics statement and patient population

This retrospective study was approved by the Local Ethics Committee for Clinical Research of the Paris Seine-Saint-Denis University Hospitals under no. CLEA-2021-185.

## 3. Results

### Sequencing quality results

Two hundred and fifty-seven clinical MTBC strains taken from patients randomly selected during routine care at Avicenne Hospital between 2015 and 2021 were sequenced. Thirty strains were excluded due to a poor quality of sequencing results: read depth at a position less than 20X.

In total, 227 strains were available for analysis, isolated from 221 patients. Sequencing quality results obtained using the PhyResSE tool showed a high average coverage of 98% against the H37Rv reference strain and an average depth of 119X. These data are not specified with Mykrobe tool.

### Comparison of WGS and phenotypic DST results

By DST, as described in Table 1, resistance to INH, RMP, EMB and PZA were 7.9%, 2.2%, 0.45% and 5.2% respectively. A total of 30 strains (13.2%) showed resistance to at least one of the first-line anti-TB drugs. Four strains (1.7%) were MDR. Regarding analysis by WGS, firstly using PhyResSE tool, we found 15 strains with a resistance to INH (6.6%), 5 to RMP (2.2%), 3 to EMB (1.3%), 11 to PZA (4.8%). Globally, resistance to at least one of the first-line anti-TB drugs was found for 20 strains (8.8%). Discrepancies between DST and PhyResSE were found for 12 strains (5.3%) detailed in Table 2. Concerning the 5 (2.2%) strains found resistant by WGS but susceptible by DST, the mutations found concerned the *embA* gene *(D4N)* for one strain, the *embB* gene *(G406A)* for one strain and the *pncA* gene *(T87M)* for 3 strains. Concerning the 7 (3%) strains found resistant by DST but susceptible by WGS, 3 strains were resistant to INH and 4 to PZA (Table 2). Secondly, using Mykrobe tool, we found 15 strains with a resistance to INH (6.6%), 6 to RMP (2.6%), 2 to EMB (0.8%), 8 to PZA (3.5%). Resistance to at least one of the first-line anti-TB drugs was found for 23 strains (10%). Discrepancies between phenotypic and genotypic methods were found for 11 strains (4.8%) for Mykrobe, detailed in Table 3. Concerning the 3 (1.3%) strains found resistant by WGS but susceptible by DST, the mutations found concerned the *rpoB* gene *(D545E)* for 2 strains and the *embB* gene *(G406A)* for one strain. For the 8 (3.5%) strains found resistant by DST but susceptible by WGS, one was resistant to RIF, 3 to INH and 4 to PZA (Table 3). The global concordance of each tool in comparison with the phenotypic DST method for determining susceptibility to first-line anti-TB drugs was 95%. The sensitivity and specificity of each tool, for the first anti-TB drugs compared to the phenotypic DST method was 70% and 98% for Mykrobe and 76% and 97% for PhyResSE (Table 1). Regarding resistance to second-line drugs, 5.3% resistance rate was detected with Mykrobe as fluoroquinolone resistance was detected in 5 strains (*gyrA* gene) and streptomycin resistance was detected in 8 strains (*rpsl* in 7 strains and *rrs* in 1 strains). A resistance rate to second-line drugs of 5.7% was obtained using the PhyResSE tool. More precisely, fluoroquinolone resistance was detected in 5 strains (*gyrA* gene), capreomycin resistance was detected in one strain (*tylA* gene) and streptomycin resistance was detected in 8 strains (*rpsl* in 4 strains and *rrs* in 4 strains).

**Table 1.**
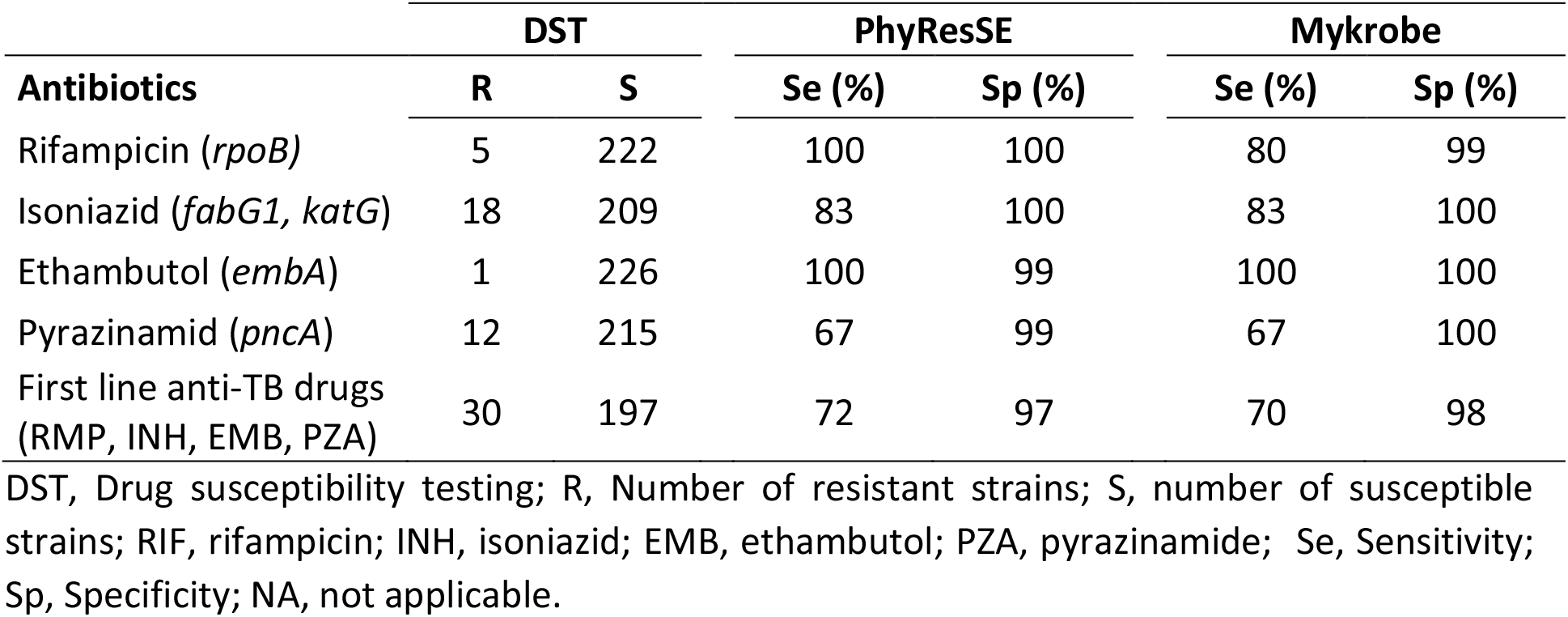
Sensitivity and Specificity of PhyResSE and Mykrobe tools compared to the phenotypic method.

**Table 2.** Phenotype-genotype mismatch identified by the PhyResSE tool.

| Drug | Phenotypically resistant |  | Phenotypically sensitive |  |  |
| --- | --- | --- | --- | --- | --- |
|  | Genetically resistant | Genetically sensitive | Genetically sensitive | Genetically resistant | Non-detected resistance |
| RIF | 5 | 0 | 222 | 0 | NA |
| INH | 15 | 3 | 209 | 0 | NA |
| EMB | 1 | 0 | 224 | 2 | One strain *embA* (D4N)<br>One strain *embB* (G406A) |
| PZA | 8 | 4 | 212 | 3 | *pncA* for the 3 strains (T87M) |

RIF, rifampicin; INH, isoniazid; EMB, ethambutol; PZA, pyrazinamide; NA, not applicable.

**Table 3.** Phenotype-genotype mismatch identified by the Mykrobe tool.

| Drug | Phenotypically resistant |  | Phenotypically sensitive |  |  |
| --- | --- | --- | --- | --- | --- |
|  | Genetically resistant | Genetically sensitive | Genetically sensitive | Genetically resistant | Non-detected resistance |
| RIF | 4 | 1 | 220 | 2 | *rpoB* for the two strains (D545E) |
| INH | 15 | 3 | 209 | 0 | NA |
| EMB | 1 | 0 | 225 | 1 | *embB* (G406A) |
| PZA | 8 | 4 | 215 | 0 | NA |

RIF, rifampicin; INH, isoniazid; EMB, ethambutol; PZA, pyrazinamide; NA, not applicable.

### Lineage determination

All the main lineages described, as the exception of the lineage 7, were represented in our dataset (Table 4). For the two tools, Lineage 4 was the most common lineage, followed by lineage 3 and 1. The two tools showed 99.5% agreement in lineage determination. Only one discrepancy was found as a strain belonging to the Euro-American lineage was identified by PhyResSE while Mykrobe identified this strain belonging to the East African Indian lineage (Table 4).

**Table 4.** Distribution of the different lineages of *Mycobacterium tuberculosis complex* strains found by the two bioinformatics tools PhyResSE and Mykrobe.

| Lineage | PhyResSE | Mykrobe |
| --- | --- | --- |
| *Mtb* Euro-American (lineage 4) | 155 (68%) | 154 (67,8%) |
| *Mtb* East African Indian (lineage 3) | 13 (5,7%) | 14 (6,1%) |
| *Mtb* Indo-oceanic (lineage 1) | 31 (13,6%) | 31 (13,6%) |
| *Mtb* East Asian (lineage 2) | 12 (5,3%) | 12 (5,3%) |
| *Mycobacterium africanum* West African 2 (lineage 6) | 6 (2,6%) | 6 (2,6%) |
| *Mycobacterium bovis* | 6 (2,6%) | 6 (2,6%) |
| *Mycobacterium africanum* West African 1 (lineage 5) | 4 (1,7%) | 4 (1,7%) |
| Total | 227 | 227 |

*Mtb, Mycobacterium tuberculosis*

## 4. Discussion

In this study, we evaluated the performances of two bioinformatics tools, Mykrobe and PhyResSE, for the analysis of sequencing data and found high performances for the two bioinformatics tools as the sensitivity and specificity compared to the phenotypic method was 70% and 98% for Mykrobe and 76% and 97% for PhyResSE, as described in other studies comparing these bioinformatics tools (26,28,29). We have chosen to study these two bioinformatics tools among others (TBprofiler, MTBseq, TGS-TBcar for example) because they are the easiest to access, i.e. usable with a standard computer on a standard search engine without any particular information skills or need to download or master other software. The two tools are easy to use and accessible to people not trained in bioinformatics. Given the facility of use, we can imagine training laboratory technicians to their use when setting up routine sequencing of MTBC strains.

Regarding the handling of each tool, we appreciated the possibility to obtain sequencing information quality offered by PhyResSE. Another advantage of the PhyResSE tool was the obtention of a bibliographic reference concerning the mutation conferring a resistance when one was identified. Nevertheless, the PhyResSE tool had some disadvantages. Firstly, the number of SNPs difference between each strain was not available. Secondly, the interface was not so user-friendly as expected in terms of presentation and ease of finding key information. The use of PhyResSE was online, which can be limiting in some work situations and the sequences (fastq files) must be uploaded on the website which was time consuming. Lastly, the uploaded sequences can be used by the designers of the bioinformatics tool.

Regarding the Mykrobe tool, we appreciated the simplicity of access to the application *via* the internet as well as its being easy to use. The presentation of the results was very clear as only the essential information was given. This application was usable to all the staff of the laboratory as well as to the clinician. We also appreciated the fact that the application can be used offline once installed on the computer. It was also possible to save the results, which allows to make verifications further when needed. Contrary to PhyResSE, MTBC sequences were not shared with designers when using the tool. The Mykrobe tool had some disadvantages: firstly, the main disadvantage was that the Mykrobe tool gave no information about the quality of sequencing in terms of coverage and depth. However, the depth was given if a mutation was detected. It is also a pity not to be able to make a phylogenetic tree with the studied strains. The developers of Mykrobe are trying to improve their application and are working on a new version, named “Mykrobe Atlas”, which would allow for real-time monitoring of global TB (30). The benefits would be to compare a patient’s bacterial DNA to all the TBs in the world and to conclude to a local outbreak or the identification of a strain that was recently seen in another country or city. Until recently, it was inconceivable to work at this scale (31).

The advantage of WGS is that it represents an “all-in-one” tool that help to investigate TB transmission but also to address the question of drug susceptibility (20). Regarding drug susceptibility, we observed a discordance between DST and WGS results for 12 strains with the PhyResSE tool (5.3%) and 11 strains with the Mykrobe tool (4.8%). The overall agreement between DST and WGS was 98.7% for the PhyResSE tool and 98.8% for the Mykrobe tool which is in accordance with other studies (32). The discrepancies observed in our study between phenotypical results and genomic results concerned the four first-line anti-TB drugs. Several large-scale sequencing studies have evaluated the correlation between SNP (single nucleotide polymorphism) identification during sequencing and phenotypically demonstrated resistance, calculating the predictive sensitivities and specificities of each SNP. According to Quan et al., the sensitivity of NGS is estimated at 94.2% and the specificity at 99.4% for an overall concordance of 99.2% for first-line anti-TB drugs (33). The discrepancies between the phenotypic and genotypic methods may depend on the regular updating of the databases used by the two tools according to the progress of knowledge. Due to its cost and performances, WGS is for now insufficient to support the use of WGS as an alternative to conventional phenotype-based DST. It should be implemented in specific cases to detect resistance rapidly in particular for the clinical management of MDR-TB strains, providing considerable information when compared with current routine methods.

The phenotypic study to second line anti-TB drugs is not performed easily in routine laboratories and is performed in majority by specialized laboratories. Contrary to standard phenotypic tests conducted in hospital laboratories, WGS offers the opportunity to determine resistance to second-line anti-TB drugs easily. The detection of an antibiotic resistance to second-line anti-TB drugs by the two bioinformatics tools is an important progress especially in the case of MDR or XDR-TB strain. Indeed, the obtention of results from phenotypic DST takes a long time and delays the adaptation of the antibiotic therapy for the patient. We found that 5.7% and 5.3% of strains were resistant to second line anti-TB drugs using PhyResSE and Mykrobe respectively. In accordance with other studies (16,17) if the WGS of MTBC was routinely implemented, the delay to get results would probably be shortened to about a week, that would be interesting in the case of a MDR or XDR-TB strains to quickly adapt anti-TB treatment. Unfortunately, the use of WGS represents a non-negligible cost in terms of human and financial resources, which can be an obstacle to its implementation in the laboratory.

## 5. Conclusion

WGS is a promising and nowadays indispensable method for the study of MTBC strains. The bioinformatics tools Mykrobe and PhyResSE were easy to use, fast and efficient. The Mykrobe tool had the advantage of being offline and its interface was more user-friendly while PhyResSe indicated sequencing information quality. The use of these platforms depends on their accessibility and update. They give access to the study of MTBC sequences to personnel not trained in bioinformatics and could be useful tools for the implementation of this technique in the laboratory. The discrepancies between phenotype and genotype observed in our study regarding antibiotic resistance remain too important to completely replace the phenotypic DST method. WGS coupled with bioinformatics tools would allow a complementary approach to standard methods for the study of MTBC strains, especially for MDR and XDR TB strains.

## Acknowledgments

We would like to thank the Molecular Biology and Data Processing Plateform of the “Université Sorbonne Paris Nord” and the Federative Institute For Biomedical Research (IFRB) of the “Université Sorbonne Paris Nord”.

